# Antibacterial activity of *Actinomycetes* isolated from the soil sample of South India and polyketide synthase gene identification

**DOI:** 10.1101/396846

**Authors:** Ashu Srivastava, Vellasamy Shanmugaiah

## Abstract

The *Streptomyces* genus is well studied owing to its capacity in producing more than 70% of antibiotics. This study was undertaken to characterize *Streptomyces* strains, occurring in soils of different districts of Tamilnadu, India as well as to evaluate their potential to produce antimicrobial compounds. Samples were collected from rice rhizosphere from different district of Tamilnadu agricultural zone. In primary screening by cross streak method, *Streptomyces strains* were assessed for antibiotic production and activity against different human bacterial. Then active isolates were selected for secondary screening by agar well diffusion method. Solvent extraction method was used to identify the best crude samples which are exhibiting better antibacterial activity. Then, 16S rRNA PCR was carried out for confirmation of isolates. In primary screening among all of the isolates, 50% isolates were active against at least one of the test organisms and 21.31% strains exhibited a broad-spectrum activity against almost all of the test bacteria. The minimum inhibitory concentrations (MICs) of the ethyl acetate extracts measured. Out of 26 positive strains two of the most active isolates SVG-07-15 and TK-01-05 were taken for further studies. These results highlight the importance of *Streptomyces* isolates in antibiotic production. Together with antibacterial activity, the PKS gene based approach can be applied for efficient screening of isolated strains of pharmaceutical value and related compounds.

**Highlights:**

- Isolation of actinomycetes from soil samples collected from South India.
- Screening of isolated actinomycetes for antibacterial activity
- Details study of potent strains against human pathogens
- Identification of PKS gene from the active strains

## 1. Introduction

Microbial diseases are increasing day by day and becoming the big problem for human health. There are more than 200 known diseases which is transmitted by bacteria, fungi, viruses, prions and other microbes to human being. The emergence of drug and multidrug resistant pathogens is the biggest issues, therefore, novel antimicrobial agents from natural resources with novel mechanism of actions are required in biopharmaceutical industry. Many research work has been carried out to control the pathogens and to identify the novel antimicrobial agents. Microbes from the soil samples are the most common natural sources exhibiting the strong biological activities against various pathogens[1]. In general, microbes produces bioactive compounds which can be useful in defence mechanism. It has been reported that the secondary metabolites produced by microbes have long benefited to human health and industry. These includes pharmaceutical agents such as penicillin antibiotics, vancomycin an anticancer agent, rapamycin an immunosuppressant among more than twenty thousand biologically active microbial natural products[2]. Secondary metabolites also have an important role as nutrient acquisition, chemical communication and defence. Polyketide synthase (PKS) and non-ribosomal peptide synthetase (NRPS) are most common among them. In bacteria, the biosynthesis of polyketide compounds such as polyene, polyether type and macrolide usually require type I PKS. Antimicrobial agents are synthetic or natural substances used to destroy or prevent the growth of bacteria, viruses and other micro-organisms (antibiotics are microbial agents which only react against bacteria). These substances have played a significant role improving public health by helping to reduce the number of deaths from diseases and infections which were previously incurable or fatal. Almost all of the living organisms have the ability to produce secondary metabolites. Overall, it has been observed that unicellular bacteria, eukaryotic fungi, and filamentous actinomyces are the most frequent and versatile producers. The filamentous Actinomycetes produces over 10,000 bioactive compounds, of which almost7600 derived from *Streptomyces*[3]. This represent the largest group (45%) of bioactive microbial metabolites. Actinomycetes, Gram positive bacteria, is one of the most important bacteria having the ability to produce a wide range of biologically active secondary metabolites against microbial pathogens. More than 80% of known antibiotics isolated from Actinomycetes and are used in medicine[4]. The genus Streptomycetes is the big producer of antibiotics[5]. Hence the present study was to investigate the antimicrobial gene and secondary metabolite from *Streptomyces* sp against human pathogens especially multidrugresistance strains of *Staphylococcus aureus*.

Presented here is the isolation of Actinomycetes from different district of Tamilnadu (such as Tuticorin, Theni and Shivaganga) and their biochemical testing, antimicrobial activities against human pathogens. About 122 isolates were screened by cross streak method against human bacterial pathogens including *Escherichia coli, Staphylococcus aureus strain A and B, Bacillus species strain A and B., K. Pneumonia, S. viridians, Pseudomonas, Klebsiella, and Pseudomonas aeruginosa*. Among 122, 61 isolates (50%) were active against at least one of the test organisms and screened secondarily by agar well diffusion method. 19 out of 61 (31.14%) isolates exhibited broad spectral activity against almost all of test bacteria[6]. Among 19 isolates, isolates namely SVG-07-15 and TK-01-05 was taken for further studies based on their superior antibacterial action against pathogens. Effective isolates were identified based on morphological, physiological and biochemical characteristics. Solvent extraction was carried out for selected isolates with eight different solvents and observed for activity. Among the solvents, ethyl acetate extract produced maximum zone of inhibition (ZOI) ranging 11-20mm against *S. aureus* when compared with commercial antibiotic oxytetracycline. Also the selected isolates were screened for polyketide identified. This will be an urgent needful research in current scenario of antimicrobial chemotherapy and drug discovery from microbial origin. Further studies on the bioactive metabolites from these cultures will be useful for discovering novel compounds of clinical and agricultural use[7].

## 2. Materials and methods

### 2.1. Sampling procedure

Soil samples were collected from agricultural land with 5-10cm depth into sterile plastic bags from Tuticorin, Theni and Sivaganga district of Tamilnadu. Soil samples were air dried at room temperature.

### 2.2. Isolation of Actinomycetes from soil samples

The isolation of Actinomycetes was done by serial dilution method. One gram of soil was suspended in 9 ml of sterile double distilled water. The dilution was carried out up to 10^-5^ dilutions. Aliquots of 1000µl were taken and spread on the Actinomycetes medium and incubated at 30°C for 7-10 days. Based on the colony growth the Actinomycetes were selected and studied further using ISP2 protocol (International Streptomyces project medium No. 2)[8].

### 2.3. Morphological Characterization of isolates

Actinomycetes isolates were inoculated on ISP2 media and incubated for 5 days at 30°C. The colonies were observed under a high magnifying lens and colony morphology were noted with respect to colour and mycelium. Mycelium of two distinct part were analyzed as aerial mycelium and substrate mycelium. There colour appearance were observed and mentioned on the table 1.

**Table 1.** Morphological analysis of Actinomycetes isolates

| **Isolates** | **Growth** | **Aerial mycelium** | **Substrate mycelium** |
| --- | --- | --- | --- |
| TK-1-04 | Abundant | WHITE TO BROWN | WHITE TO YELLOW |
| TK-1-05 | Abundant | WHITE | LIGHT YELLOW |
| TK-1-03 | Good | BROWN | YELLOW |
| TK-8-02 | Good | LIGHT BROWN | BROWN TO DARK YELLOW |
| TK-8-03 | Abundant | DARK BROWN | YELLOW |
| SVG07 | Abundant | WHITE TO GREEN | DARK BROWN |
| SVG06 | Abundant | LIGHT BROWN | BROWN TO YELLOW |
| NL-5-01 | Abundant | WHITE TO BROWN | YELLOW TO DARK BROWN |
| NL-5-02 | Abundant | BROWN | BROWNISH GREEN |
| NL-5-03 | Good | WHITE TO GREEN | YELLOW |
| NL-5-04 |  | WHITE TO BROWN | YELLOW |
| NL-5-07 | Good | WHITE TO LIGHT BROWN | LIGHT BROWNING YELLOW |
| NL-5-08 | Abundant | WHITE TO LIGHT BROWN | WHITISH YELLOW |
| THN-5-01 | Abundant | GREENISH BROWN | YELLOW |
| THN-5-02 | Abundant | WHITE TO GREEN | BROWN TO DARK GREEN |
| THN-5-03 | Good | GREEN | BROWN TO DARK GREEN |
| THN-5-04 | Abundant | WHITE TO LIGHT BLUE | YELLOW TO DARK BROWN |

### 2.4. Screening for antimicrobial activity

The antimicrobial activity of isolated Actinomycetes were performed by cross streak method. The plates were prepared and inoculated with isolated by single streak at the centre of petri dish and incubated at 30°C for 3 days. The palates were then inoculated with test organism by a single streak at 90° angle to the Actinomycetes strains and incubated at 37°C overnight. Then the antagonism of test organism was recorded. The human bacterial pathogens such as *Escherichia coli, Staphylococcus aureus strain A and B, Bacillus species strain A and B., K. Pneumonia, S. viridians, Pseudomonas, Klebsiella, and Pseudomonas aeruginosa*were used. The positive isolates were further screened against these pathogens by using agar well diffusion method. The pathogens were swabbed on NA plates and incubated at 37°C for 24 hours. The wells were prepared using sterile cork borer at five positions. Different concentration (0, 25, 50, 75 and 100µl) of isolates were incubated against the pathogens to observe the antimicrobial activities[9].

### 2.5. Biochemical Testing

After preliminary studies, the isolates which were found to be positive were selected for biochemical testing. These includes: Catalase test, Indole test, Methyl red test, Voges-Proskauer test(VP test), gelatine hydrolysis, starch hydrolysis, urea hydrolysis, acid production from different sugars, hydrogen sulphide (H_2_S) production test, motility test, triple sugar iron (TSI) agar test, and citrate utilization test.

### 2.6. Genomic DNA extraction

Actinomycetes isolates were inoculated aseptically into a 250ml Erlenmeyers flask containing 30 mL of the ISP2 media. Incubate up to the log phase in a rotary shaker at 30°C at 180 rpm. Addition of glycine (20%) and MgCl_2_ into the media. Centrifuge the culture at 10,000 rpm for 10 minutes. Wash the culture twice with lysis buffer. Transfer the mycelia into fresh tube containing 500 μl of TE buffer supplemented with lysozyme (50 mg/mL). Incubate the mixture for one hour. Take 100µl of this mixture and use DNA isolation kit (Qiagen) for DNA isolation according to the manufacturer’s instruction.

### 2.7. Screening for Polyketide Synthase gene

It is well known that many bioactive metabolites in Actinomycetes are produced by PKS gene. Screening for genes associated with secondary metabolism is helpful in evaluating the biosynthetic potential of Actinomycetes. PCR was performed using the following amplification parameters: initial denaturation at 94°C for 5 min; then 35 cycles of 94°C for 1 min, annealing at 50°C for 1 min, extension at 72°C for 2 min; and a final extension at 72°C for 10 min. The amplified product were assessed by agarose gel electrophoresis. Primer used for PCR amplification were 540F, 5’-GGITGCACSTCIGGIMTSGAC-3’ and 1100R, 5’-CGATSGCICCSAGIGAGTG-3’.

### 2.8. Extraction of Bioactive Compound

Production of bioactive compounds was done by submerged fermentation. Active Actinomycetes isolates were taken in 50 ml of ISP2 broth in 250 ml capacity conical flask under sterile conditions and incubated at 30°C for 7 days at 150rpm rotation. After the fermentation, the medium was centrifuged at 10000 rpm to removed cell debris[10]. Different solvents such as; benzene, ethyl acetate, petroleum ether, chloroform, hexane, acetone, methanol and ethanol were used to extract the secondary metabolites from the isolates. Among these solvents ethyl acetate gave highest crude products. Therefore, resultant fermentation broth were added with the equal volume of ethyl acetate solvent. Then the sample were shaken vigorously in rotary shaker. The solvent phase were collected in a beaker and kept for evaporation using desiccators. The completely dried residues were re-dissolve in DMSO and used for further analysis.

### 2.9. Antibacterial activity of ethyl acetate extract against human pathogens

The most active isolates such as SVG-07-15 and TK-01-05 were further studied for their antibacterial activity. The partially purified crude extract was determined by agar well diffusion method. Bacterial concentration of *S. aureus* A and B were adjusted at 0.5McFarland turbidity standards and inoculated on Nutrient agar plates by using sterilized cotton swabs. Wells were bored by sterilized cork borer. Different concentration of 25, 50, 75 and 100µl of crude extract were poured into wells. Plates were incubated for 24 hours at 37°C.

## 3. Results & Discussion

### 3. 1. Isolation & Morphological analysis of Actinomycetes

Soil Actinomycetes were isolated from the agricultural land of different district of Tamilnadu. After 7 days of incubation the growth of Actinomycetes were observed on the plate. The number of Actinomycetes colonies in each dilution plate did not necessarily follow the 10-fold serial dilution pattern as expected, but varied across each dilution and colony type. A total of 122 isolates were grown and maintained using ISP2 medium. Fig. 1 shows the growth of Actinomycetes isolates from the four different district’s soil samples. The morphological pattern of grown Actinomycetes isolates were categorised as per their growth pattern, substrate and aerial mycelium (Table 1). Isolates growth was either in good or abundant condition. The substrate and aerial mycelium shows the colour appearance of individual isolates which can be either white/brown or some changes in these colour appearance.

**Fig. 1.**
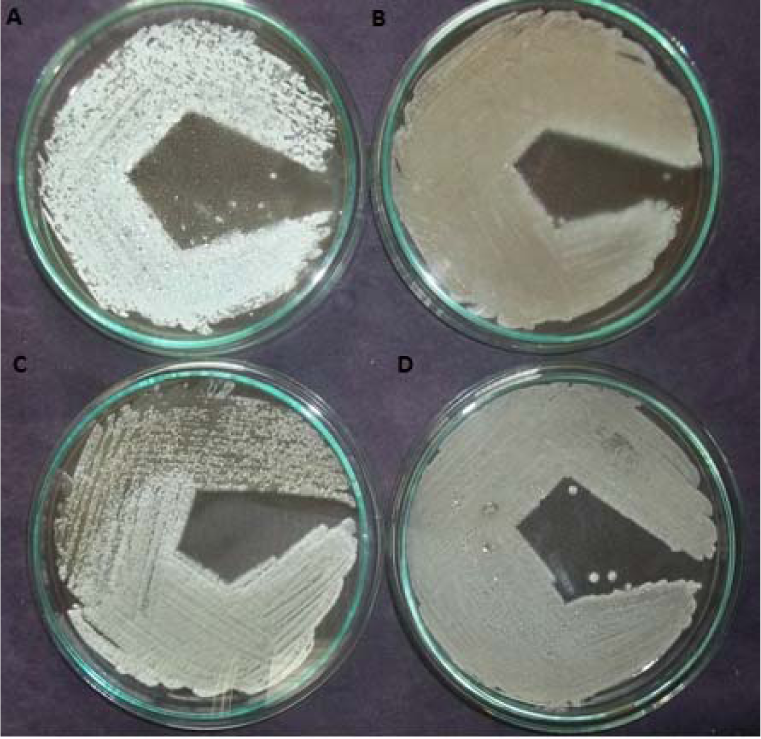
Actinomycetes isolates on ISP2 medium, A. Shivganga (SVG), B. Nellai(NL), C. Tuticorin (TK), and D. Theni (THN).

### 3. 2. Antimicrobial Screening

#### 3. 2. a. Preliminary screening

Cross streak method were used to calculate the minimum inhibition concentration (MIC) value of isolates against different human pathogens. The test pathogens were *Escherichia coli, Staphylococcus aureus strain A and B, Bacillus species strain A and B., K. Pneumonia, S. viridians, Pseudomonas, Klebsiella, and Pseudomonas aeruginosa.*The Actinomycetes isolates grown at the centre of the plates and further perpendicularly the pathogen strain were incubated for 24 hours to observe their antibacterial effect. The control plates without the Actinomycetes isolates were also prepared to compare the similar pattern of growth (Fig. 2). The MIC value were measured in mm. Among 121 isolates of Actinomycetes there was a total of 61 isolates shows antibacterial effect against one or more pathogens. The histogram were plotted which shows the positive isolates SVG-07-15 and TK-01-05 against test pathogens performed in triplicate (Graph 1).

**Fig. 2.**
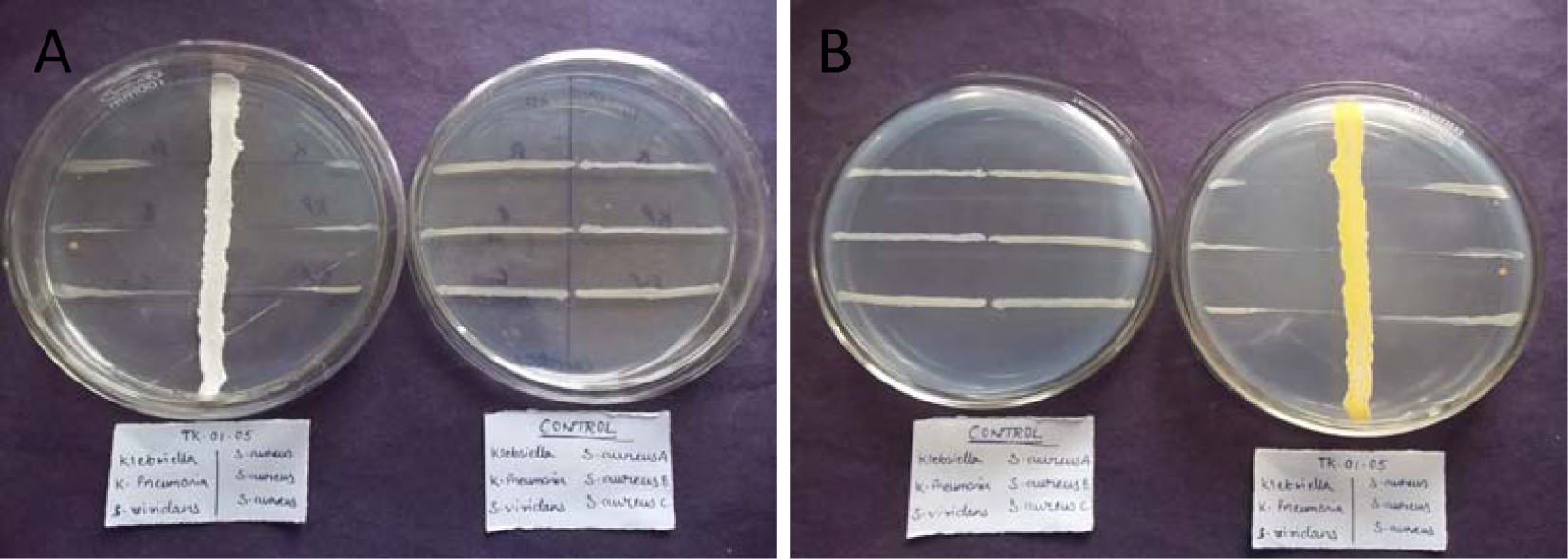
Cross streak method of Actinomycetes against human pathogen test vs. control A. Front, B. Backview.

**Graph 1.**
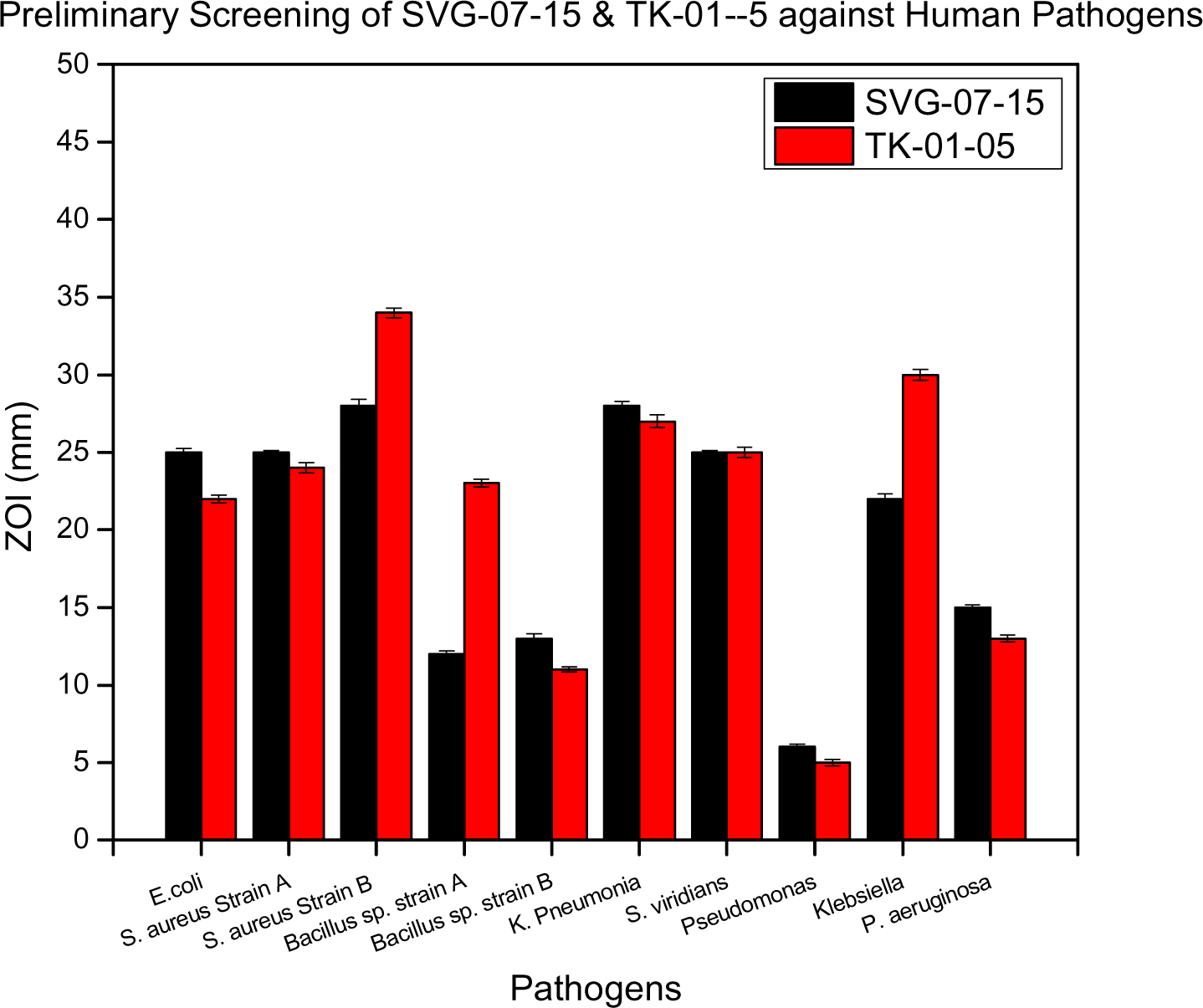
A histogram for the two positive isolates SVG-07-15 and TK-01-05 against test pathogens

The two most potent isolates represents the antibacterial property against human pathogens. The zone of inhibition is maximum in case of the pathogen *S. aureus* B and A strains while the minimum against Pseudomonas. This indicates that the potential for these two strains against *S. aureus* can lead to the further investigation towards multi-drug resistant *staphylococcus aureus*(MRSA).

#### 3. 2. b. Secondary screening

Agar well diffusion method were used to calculate the zone of inhibition (ZOI) against the pathogens. Isolates selected from the preliminary screening were further utilized. Different concentrations (25, 50, 75, 100 µl) of selected isolates shows different zone of inhibition (Fig. 3). Lowest concentration (25 µl) having small zone of inhibition while highest concentration (100 µl) having biggest zone of inhibition. As the concentration increases the ZOI also increases. Among 61 isolates a total of 25 isolates shows antibacterial effect against these pathogens.

**Fig. 3.**
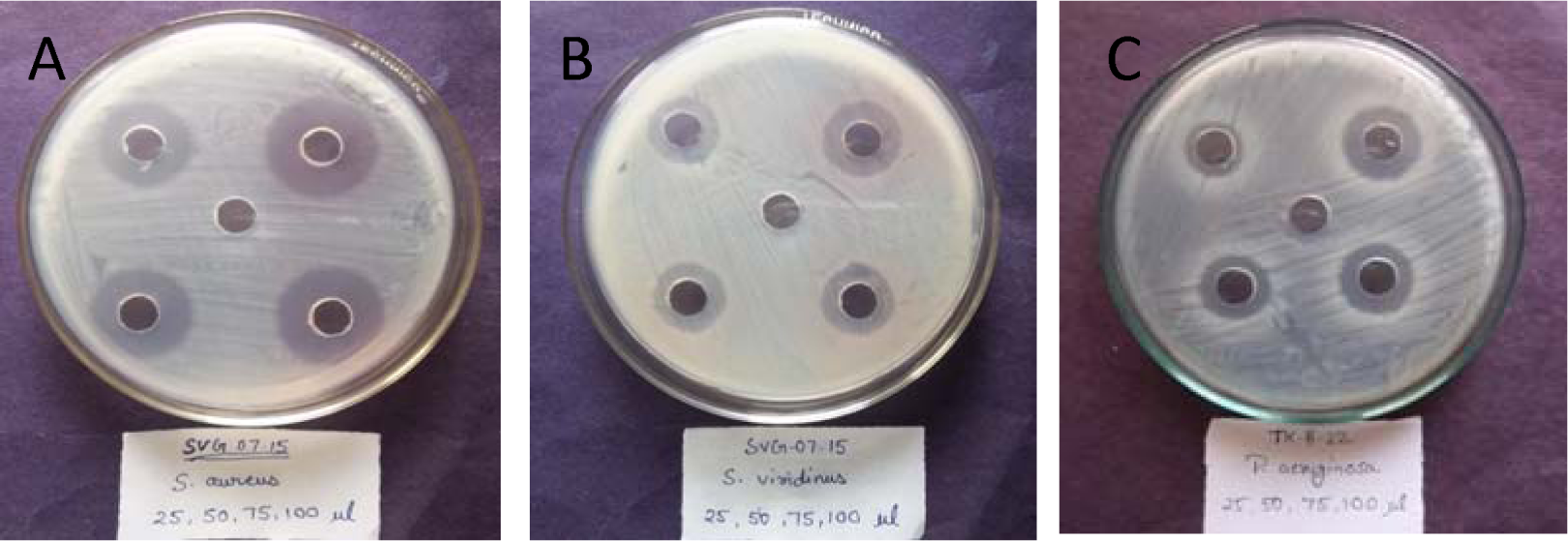
Agar well diffusion method for the Actinomycetes isolates against A. *S. aureus,* B. *S. Viridinus and C. P. Aeruginosa.* The Zone of inhibition were measured.

### 3. 3. Biochemical analysis

Different biochemical testof the selected 25 isolates were performed (Fig. 4). These test results confirmed about the properties ofActinomycetes. Catalase test facilitate the detection of enzyme catalase in bacteria. The positive reaction are evident by immediate bubble formation. No bubble formation represents a catalase negative reaction. Gelatin test gives the result of either strong or weak positive i.e. liquefaction occurs within 3-4 days or negative which means no liquefaction even after 30 days. Indole test reagents are used to detect the production of indole by bacteria growing on media containing tryptophan. In case of MR test the culture medium turns red after addition of methyl red, because of a pH at or below 4.4 from the fermentation of glucose, the culture has a positive result for the MR test. While negative results indicated by a yellow colour in the culture medium, which occurs when less acid is produced (pH is higher) from the fermentation of glucose. Starch hydrolysis test shows the production of amylase. TSI test a microorganism’s ability to ferment sugars and to produce hydrogen sulfide. Bacteria that ferment any of the three sugars in the medium will produce by products. These by products are usually acids which will change the colour of the dye.

**Fig. 4.**
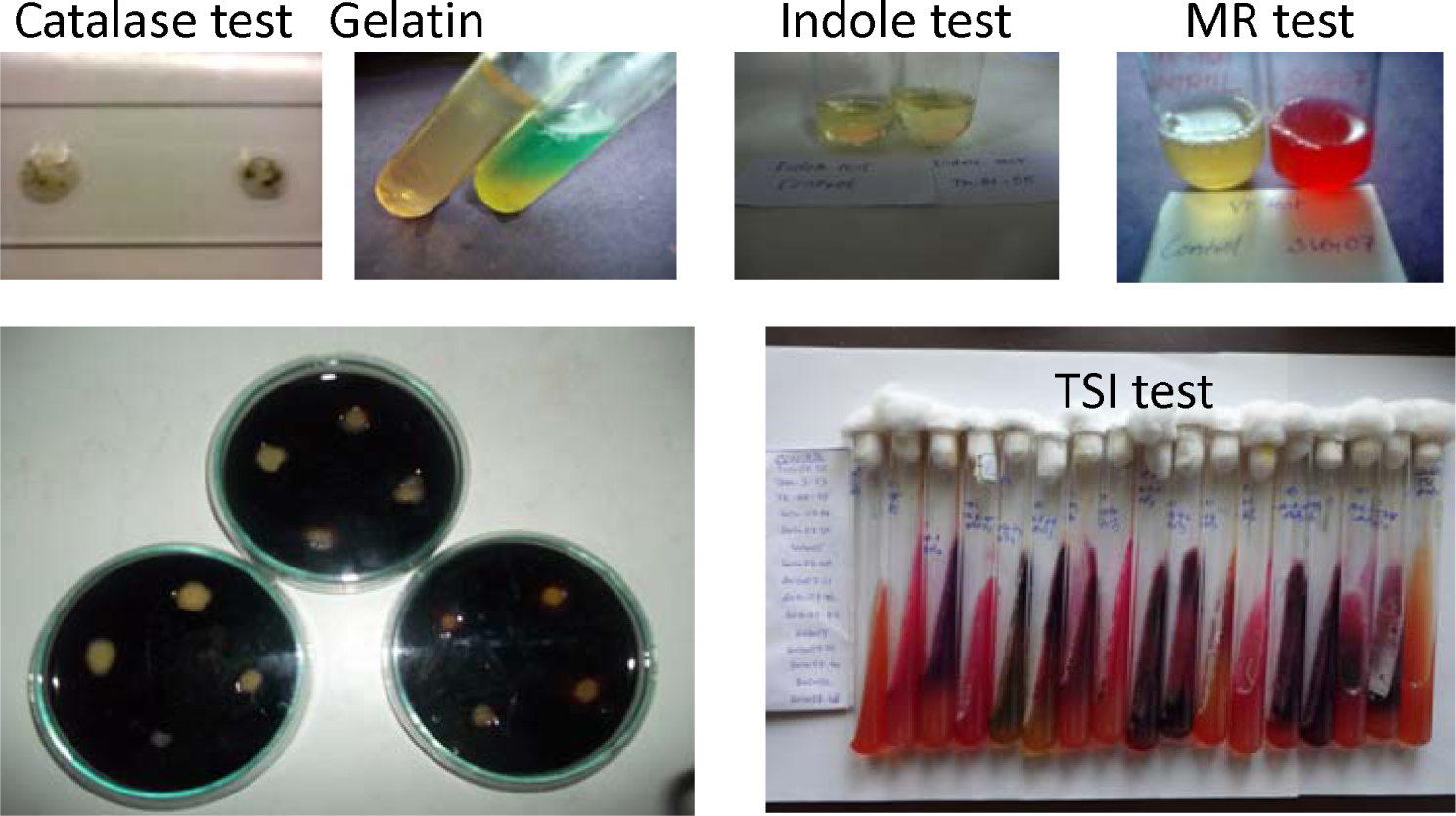
Biochemical test of Actinomycetes isolates such as Catalase test, Gelatin liquefaction test, Indole test, Methyl red test, Starch hydrolysis test and triple sugar test.

### 3. 4. Genomic DNA &Amplification for Polyketide Synthase gene

Genomic DNA were isolated from the screened Actinomycetes isolates and run on agarose gel electrophoresis. Four positive isolates of Actinomycetes were selected for the amplification of PKS gene from a total of 25 positive strains obtained by secondary screening.PCR amplification of the PKS I gene which were expected yield was 1400bp were performed and run on agarose gel electrophoresis (Fig. 5). The isolates shows a prominent band on the gel against the 1 KB ladder. From the preliminary morphological studies it was observed that the isolates belong to Streptomyces spp. The richest group of Actinomycetes represented by the Streptomyces which has been extensively isolated by exploring antibacterial, antifungal and other biologically active substances. Therefore, these screened positive isolates were amplified on PKS genes and all of these revealed that strains had gene expression for biosynthetic of polyketide substances. The PCR primer used to detect the presence of PKS genes in the strain.

**Fig. 5.**
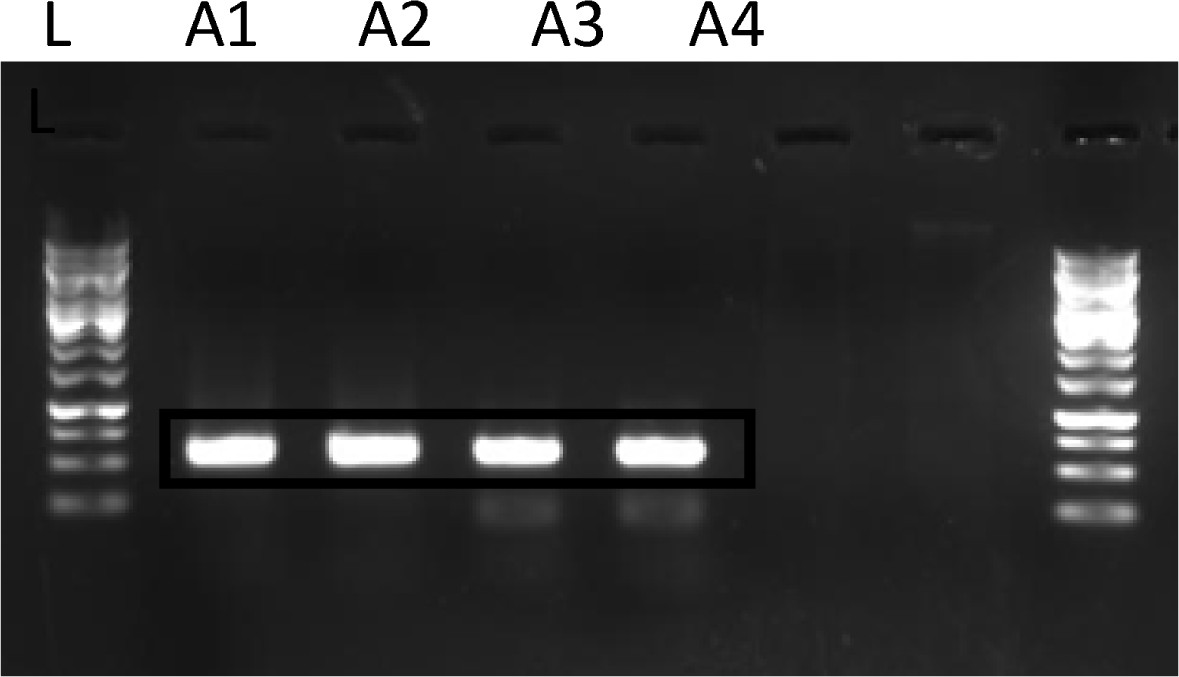
PCR amplification for the PKS I gene identification from Actinomycetesisolates where L is the ladder, A1 is TK-01-04, A2 is TK-01-05, A3 is TK-08-22, and A4 is SVG-07-05.

### 3. 5. Antibacterial activity of crude against human pathogens

Crude extract of the isolates SVG-07-15, TK-01-05, TK-01-04 and TK08-22 have been shown to have significant effect against human pathogens such as *S. aureus strain B*(Fig. 6). The statistically significant zone of inhibition were measured and compared with standard tetracycline and penicillin which is tabulated below (Table 2). It has been observed that as the concentration of crude product increases in the well, the zone of inhibition also increases. This represents the concentration dependant activity of crude extract. Crude extract from isolate SVG-07-15 showed high antimicrobial activity against *S.aureus* in comparison with standard amoxicillin and tetracycline. Even the isolate TK-01-05 also shows the higher activity with increasing concentration against both the strains A and B of *S. aureus*.

**Fig. 6.**
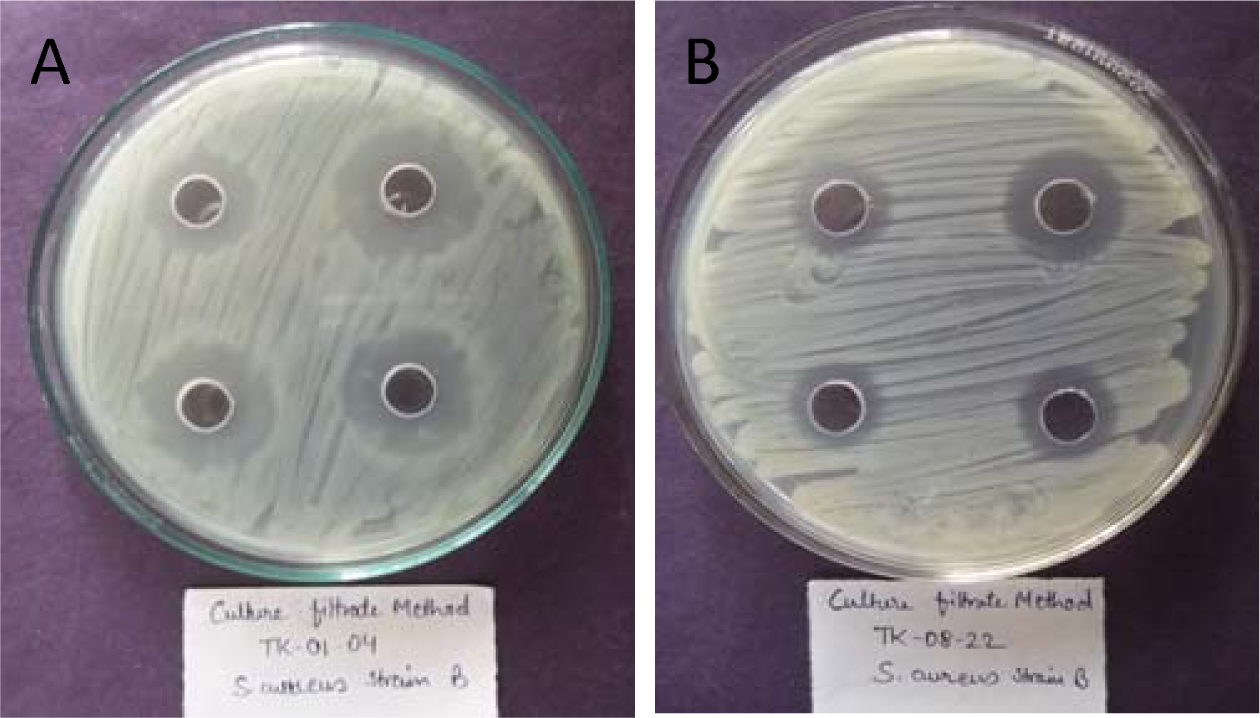
Culture filtrate method using crude extract of ethyl acetate with different concentration for the Actinomycetes isolates against *S. aureus*.

**Table 2.** Secondary metabolites produced using ethyl acetate crude extract were measured for the zone of inhibition against *S. aureus.* Values are mean±SD

| Isolates | Zone of Inhibition (mm) |  |  |  |  |  |  |  |
| --- | --- | --- | --- | --- | --- | --- | --- | --- |
|  | *S. aureus* A |  |  |  | *S. aureus* B |  |  |  |
|  | 25µl | 50µl | 75µl | 100µl | 25µl | 50µl | 75µl | 100µl |
| **SVG-07-15** | 16±1 | 18±1 | 20 | 20±1 | 12±0.5 | 16 | 18±1.5 | 19±0.5 |
| **TK-01-04** | Nil | 11±2 | 12±1.5 | 13±0.5 | Nil | Nil | 13±2 | 13±1 |
| **TK-01-05** | 14±1.5 | 16 | 17±1 | 20±2 | 12±1.5 | 14±0.5 | 14 | 16±2 |
| **TK-08-22** | 08±2 | 10±1 | 11±0.5 | 14 | 06 | 08±1.5 | 09±1 | 11 |
| **Tetracyclin** | 32±1 | 32±1 | 32±1 | 32±1 | 31±1 | 31±1 | 31±1 | 31±1 |
| **Penicillin** | 0±1 | 0±1 | 0±1 | 0±1 | 0±1 | 0±1 | 0±1 | 0±1 |

## Conclusion

The screening of soil bacteria for novel bioactive compounds shows great attention in research. The isolated strains having antibacterial, antifungal activity, which tends to enrich compounds that are already known and abundantly present in environment. Around 23,000 bioactive secondary metabolites produced by the different microorganisms almost 10,000 of these compounds are produced by Actinomycetes[11]. A wide range of antibiotics in the market obtained from Actinomycetes. Presented data also shows the effect of soil isolates against different human pathogens. The most positive strains were deeply studied and their secondary metabolites were produced using solvent. Even the identification of polyketide genes were performed to know their possible antibacterial activity. In summary combined with antibacterial activity and the PKS gene based molecular approach can be applied to efficient screening of strains with pharmaceutical values. Thus our study can help in further depth of knowledge regarding the environmentally present soil sample[12]. For novel drug delivery, researchers still exploiting the chemical and biological diversity from diverse Actinomycetes group to maximize the possibility of successful discovery of isolates in cost effective manner.

## Acknowledgements

Authors are thankful to the DBT-IPLS for the financial support. We are grateful to Microbial Gene Technology department of MKU to perform the entire experiment.

